# An algorithm that combines machine learning ensemble modeling and network analysis to predict self-tolerant tumor-associated antigens for anti-cancer immunotherapy

**DOI:** 10.1101/2023.02.09.527464

**Authors:** Christopher Lischer, Martin Eberhardt, Adrian Weich, Julio Vera

## Abstract

Tumor-associated antigens (TAAs) and their derived peptides constitute the chance to design off-the-shelf mainline or adjuvant anti-cancer immunotherapies for a broad array of patients. Here, we present a computational pipeline that selects and ranks candidate antigens in a multi-pronged approach and applied it to the case of uveal melanoma. In addition to antigen expression in the tumor target and in healthy tissues, we incorporated a network analysis-derived antigen indispensability index motivated by computational modeling results, and candidate immunogenicity predictions from a machine learning ensemble model on peptide physicochemical characteristics.

## Introduction

Uveal melanoma (UM) is the most frequent primary ocular malignancy in adults, with an incidence of 5.1 per million that has remained stable in the US over the past five decades (PubMed identifier, PMID: 15651058). In general, overall survival (OS) times of UM have not improved over the past decades, emphasizing the need for alternative treatment options (PMID: 32273508).

Recent research effort in different cancer entities has focused on targeting tumor-associated antigens (TAA) with different strategies like therapeutic mRNA vaccines encoding said antigens (PMID: 32244193) or direct vaccination with engineered peptides (PMID: 30524907), These approaches ultimately stimulate the adaptive immune response against the targeted antigen by engaging major histocompatibility complex class I (MHC-I) interactions with activated cytotoxic T lymphocytes (CTL, PMID: 28367149).

Finding antigens that are well-suited for therapeutic immunotherapy has proven to be a challenging and up to now unrewarding undertaking. Our goal in this study was to develop a computational predictor for safe TAAs. Thus, we integrated models addressing the above complications into a practical and deployable algorithm. In an ensemble model approach, we combined multiple data sources and estimates to predict the treatment efficacy of TAAs and their derived epitopes. We provide the annotated results as a database publicly and free for non-commercial use at https://www.curatopes.com/uvealmelanoma.

## Methods

### Prioritization of genes from transcriptomics data

The underlying algorithm to prioritize genes was described in detail in a previous publication (PMID: 31416842). Briefly, genes needed to meet four criteria: 1) annotated as protein-coding, 2) no histological evidence of protein expression in normal tissue according to the Human Protein Atlas (HPA, proteinatlas.org, PMID: 25613900), 3) RNA-seq-based expression of above 1 TPM in at least 90% of UM samples, 4) RNA-seq-based expression in 90% of normal tissues according to GTEx (stratified by tissue, PMID: 26484571) lower than in 90% of UM samples. In sum, these combined the eligible-protein with the favorable-transcript set as defined previously.

### Network analysis and gene indispensability index

To reduce the risk of antigen loss rendering the targeted therapy ineffective, we implemented a network-based method to quantify a gene’s indispensability for tumor growth and survival. In a first step, we calculated for each gene its inherent importance by adding the number of its associations with a manually curated list of 90 cancer-relevant GO terms **(Suppl. Table S1)** and the number of its occurrences in the four databases Oncogene (PMID: 28162959), the Cancer Proteomics Database (http://apoptoproteomics.uio.no/), the Epithelial-Mesenchymal Transition Gene Database (PMID: 31941584), and DriverDBv3 (PMID: 31701128). We performed this calculation for our prioritized genes and all genes annotated in DriverDBv3 to obtain a cancer entity-agnostic background distribution of gene importance.

To incorporate the fact that proteins execute their functions embedded in biochemical networks, a gene’s indispensability index was designed to be higher when its loss disrupts close interactions with other genes of high importance. In a second step, we therefore automatically reconstructed an interaction network from our prioritized genes (PMID: 28137890), and all genes annotated in DriverDBv3 and expanded it with direct interactors extracted from the databases TRANSFAC, HTRIdb, miRecords, and miRTarBase. Each network node was then assigned a neighborhood importance (NI) calculated as the sum of its own and its direct interactors’ gene importance. The higher a gene’s NI, the more relevant it is for cancer development. See **Supplementary Figure S1** for further details on the calculation.

We conjectured that there is no real gain of importance above a certain threshold value of NI, or conversely that a gene whose NI equals the threshold and another gene with NI above threshold represent, in our context, almost identical liabilities for a cancer cell. Therefore, we decided to transform the NI’s empirical distribution into one with saturation characteristics. We defined the aforementioned threshold as 90% of the maximum in a saturation kinetics function of Michaelis-Menten type. The threshold’s numerical value *t* was set by multiplying two cutoffs derived from the empirical distributions of two variables related to NI, i.e., node degree and gene importance: first, the node degree cutoff, as a measure for “sufficiently highly connected”, was set to 5, and then the gene importance cutoff, as a measure for “sufficiently important gene”, was found by calculating the empirical cumulative probability *p* of the value 5 in the node degree distribution and selecting the gene importance’s corresponding p-quantile. In this manner, the two cutoffs effectively mark the same upper-end fraction of their respective distributions as saturated.

After plugging the threshold value *t* and a saturation level of 90% into the Michaelis-Menten equation and reordering, the saturation kinetics’ K_M_ parameter was calculated according to equation I.

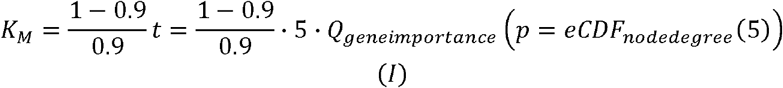

The transformed neighborhood importance of a gene G according to equation II, displaying saturation behavior and confined to the unit interval, was labeled indispensability index.

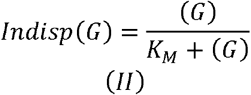

### Generalized binding and activity prediction using machine learning

We enumerated MHC-I-restricted peptides from the Ensembl-derived canonical FASTA protein sequences corresponding to the prioritized genes. To rule out autoimmune reactions due to coincidental sequence identity, we compared the enumerated peptides against all non-prioritized genes in the human proteome and discarded peptides for which we found an exact sequence match. For the remaining peptide set, we predicted their allele-specific MHC binding affinities with netMHCpan v4.0 for all 36 HLA alleles available in this version (PMID: 32406916, **Suppl. Table S2).**To complement these with an allele-independent measure of peptide immunogenicity, we constructed two random forest (RF) machine learning models to predict which peptides have a high chance of generalized MHC-I binding (gBP) or of eliciting an immune response (gAP).

The models were implemented in R with the library randomForest and designed to accept the following physiochemical peptide features as input: hydrophobicity, isoelectrical point, molecular weight, stability index, polarity, and sequence length. Apart from polarity, whose derivation relied on our own code founded on literature, the features were predicted with the Biopython (PMID: 19304878) module ‘ProtParam’. Training MHC-I-restricted peptides for both models were extracted from the MHCBN database (PMID: 19379493) 4.0 by selecting only peptides with unambiguous annotation (i.e., yes or no) for binding or activity, respectively. This yielded 3610 binders vs 368 nonbinders, and 788 active vs 475 non-active peptides. The training set for binding was supplemented with 101 binders and 100 non-binders identified through crystallography experiments.

To more closely model the expected distribution of binders to non-binders in empirical data, we performed weighted sub-samplings of the relevant training data by constructing the input set in such a manner that the ratio of binders to non-binders was 1:10. Since this in combination with the set-wide binder-to-non-binder ratio of roughly 10:1 reduced the effective input set size considerably, we performed 100 iterations of weighted sub-sampling from the training data and for each trained an RF model with 10,000 trees. For the activity probability, we used a balanced split (active to not active, 1:1) because, in theory, any peptide can elicit an immune response when engaging a complementary T-cell receptor. For both models, responses were discretized at the threshold of 0.5 and predictive power evaluation was performed against the entirety of the input datasets. The respective averages of the 100 RF models’ discretized classifications were used as the probability output of gBP and gAP.

### Derivation of peptide efficacy score

The efficacy score ES of an epitope results from a multi-criteria function that aggregates five components derived in the above computations. Each individual component was constrained to the range between 0 and 1 to ensure unbiased contribution. The full formula is

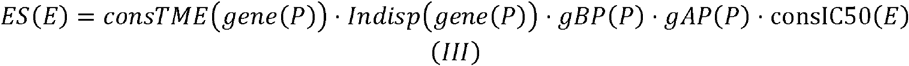

with *P* denoting a peptide, gene(P) denoting its corresponding gene of origin, and E denoting a full epitope, i.e., a combination of *P* with a specific HLA allele. The components and outputs are efficacy score ES, constrained tumor median expression consTME, indispensability index Indisp, generalized binding predictor gBP, generalized activity predictor gAP, and constrained binding affinity consIC50. Constrained in the cases of consTME and consIC50 means that we transformed the observed value range to the unit interval [0,1] with piece-wise rules as follows:

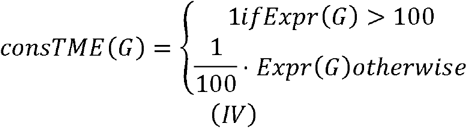

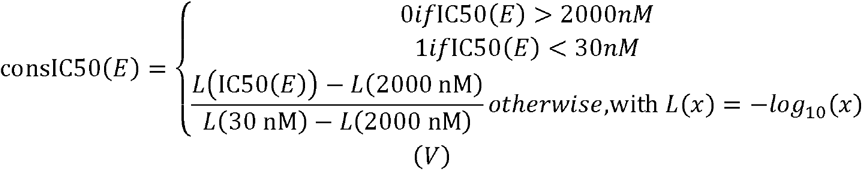

with *G* and *E* denoting gene(P) and epitope(P), respectively, Expr(G) denoting G’s RNA expression, and IC50(*E*) denoting the binding affinity predicted with netMHCpan. The upper bound for gene expression (100 TPM) was selected according to the rationale utilized in Lischer *et al*. The binding affinity bounds, 2000 nM and 30 nM, respectively, were derived from a logistic regression applied to the training dataset of the RF models above and were optimized for a high positive predictive value to reliably discard non-binders (see further discussion in Supplementary Material).

### Selection of peptide candidates for validation

For validation, we selected three distinct tiers from our pipeline output – high efficacy (HE), low efficacy (LE), and alternative predictor (AP).

The high-efficacy (HE) tier contained the top-ranked peptides we deemed well-suited for therapy. To maximize donor availability for experimental validation, we selected peptides with high efficacy scores for the locally prevalent HLA allele A*02:01 (abbreviated A2) as follows: For each scored peptide, we first assessed its potential to engage bystander alleles, i.e., any of the 35 considered alleles beyond A2 **(Suppl. Table 2),**which would wrongfully inflate the observed experimental signal due to non-A2-mediated stimulation. The probability of engaging at least one bystander allele was estimated by interpreting a peptide’s bystander efficacy scores as probabilities of success and calculating the probability of at least one success across all bystander alleles, i.e., the non-failure probability *P_NF_*:

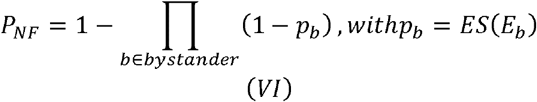

In addition, the maximum of the bystander efficacy scores per peptide were calculated. Peptides were then ranked by multisorting their attributes in the following order: bystander non-failure probability (ascending), maximum of bystander efficacy scores (ascending), and A2 efficacy score (descending). This multisort step gives priority to discarding peptides with undesirable binding to bystander HLA alleles. In a subsequent step coined Levenshtein filter, we ensured that no two peptides in the final selection are highly similar. For this, we sequentially discarded peptides whose sequences differed by only one amino acid substitution, insertion, or deletion (i.e., showed a Levenshtein string distance of 1) from a peptide of higher rank. The top 20 peptides from the remaining list made up the HE tiers.

Low-efficacy (LE) peptides were those with minimal efficacy scores across all 36 considered HLA alleles. To ensure a randomized order of the large number of peptides with efficacy scores of 0 across all alleles, peptide order was shuffled before sorting. Then, for each peptide, the product, maximum, and mean of its scores across all 36 alleles were calculated and the amount of its non-zero scores counted. A subsequent all-ascending multisort on these four features in the aforementioned order ranked the peptides in order of increasing efficacy profile. After applying a Levenshtein filter as described above, the top twenty peptides were selected for the LE tier.

The 20 peptides in the alternative-predictor (AP) tier served as theoretically efficacious counterparts to the HE peptides to examine how our selection pipeline performs in comparison to established methods. They were chosen from the set of peptides assigned an A2 efficacy score of zero. After application of the Levenshtein filter, the AP tier was filled by selecting the 20 peptides with the closest marginally better IC50 value predictions to the HE tiers.

## Results

In this work, we construct and deploy a methodology to rank the therapeutic efficacy of potential antigenic peptides for a specific tumor of interest (**Figure 1**). Apart from properties relating to the antigen’s efficacy in stimulating an immune response, we gauge safety in other tissues and risk of immune evasion, and finally check the plausibility of our predictions in *in vitro* experiments.

**Figure 1.**
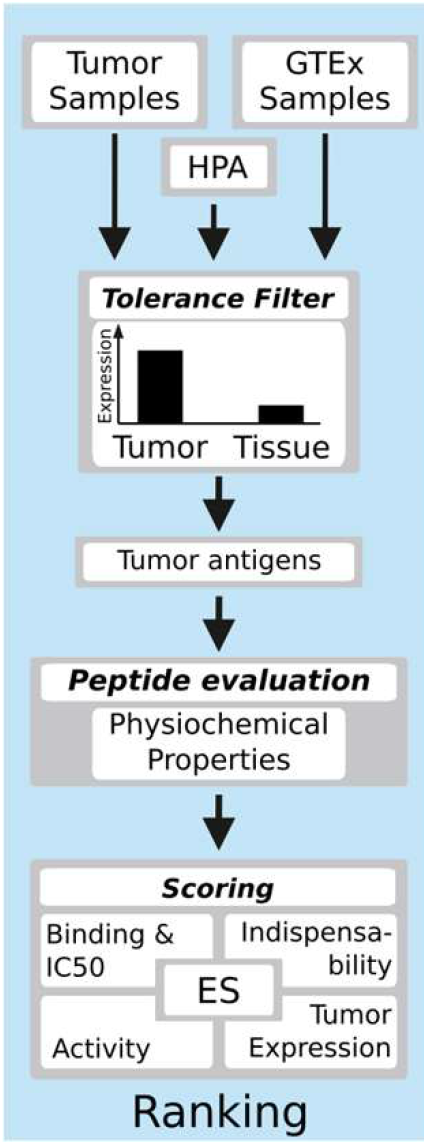
Overview of the algorithm. We evaluated and filtered genes based on their expression profiles to create a database of tumor antigens that we propose as optimized candidates for targeted anti-cancer therapy.

### Gene prioritization and peptide evaluation

In the first step, we applied the first phase of the algorithm described in Lischer *et al*. (PMID: 31416842) to identify protein-coding genes with a high-in-tumor, low-in-tissue expression profile from the empirical mRNA abundancies of 80 primary uveal melanoma (UM) samples (Robertson, PMID: 28810145) (see Method section for details). Twenty-two genes were selected (CCDC140, TRAPPC9, C14orf169, C11orf71, ACCSL, SMIM10L1, MLANA, PMEL, TYRP1, TSPAN10, RAB38, ABCB5, CABLES1, OCA1, TRPM1, SLC45A. FNDC10, TMEM200C, PNMA6A, ALX1, ELFN1).

To estimate a gene’s indispensability, we extracted from GO terms and databases the amount of its associations that benefit tumor survival. Projecting these numbers onto a reconstructed UM-tailored signal interaction network, we derived a normalized gene-specific indispensability index **(Fig. 2B** and **Suppl. Figure 1**) that was used in the later ranking of epitopes.

**Figure 2.**
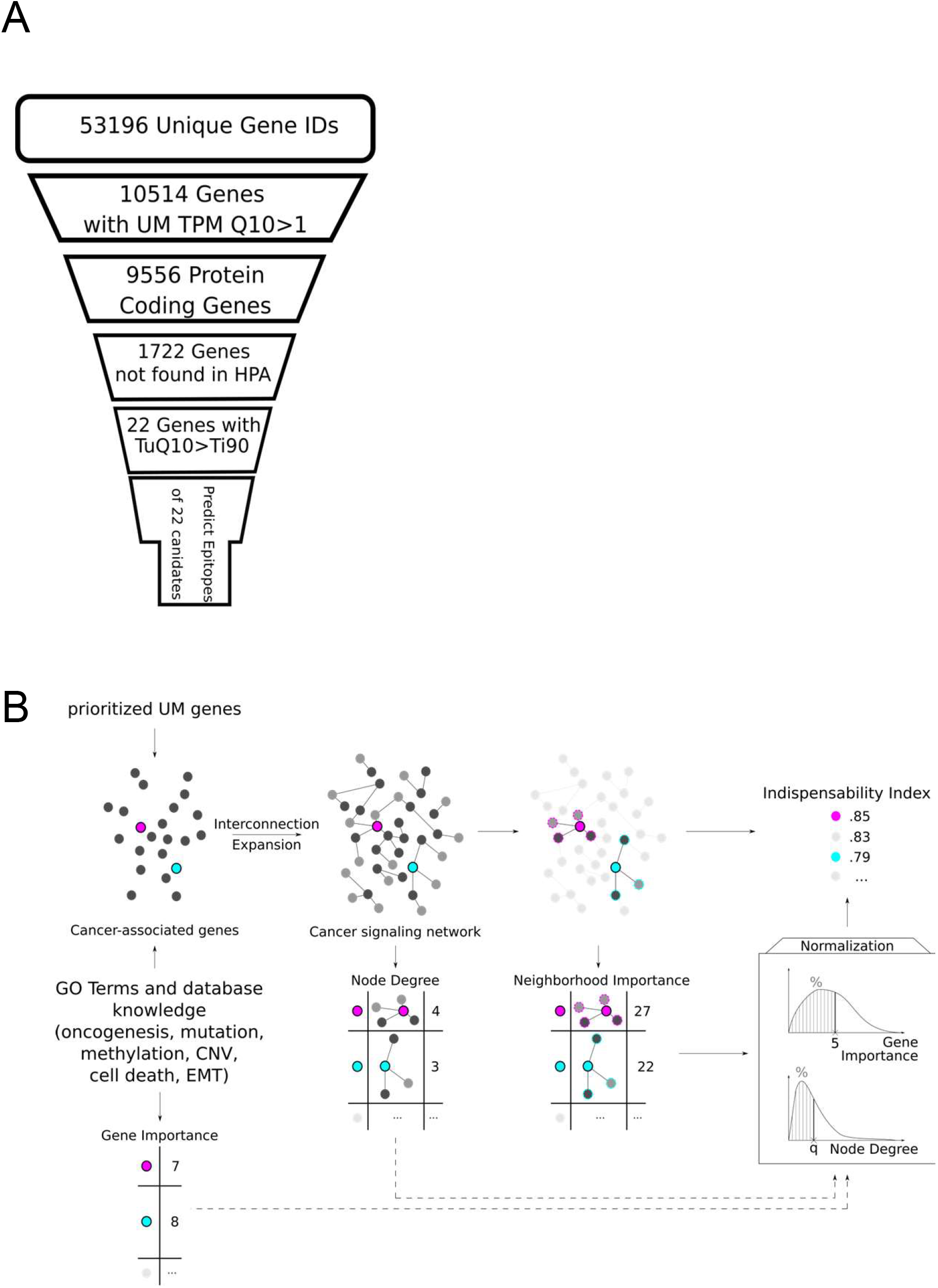
Selection and network evaluation of candidate genes. **(A)**Selection funnel representing a cascade of *in-silico* filters for genes. Each slice of the funnel lists the feature criterion and the number of genes meeting it. Tumor expression statistics were calculated based on a published set of 80 primary UM samples. Ultimately, 22 candidate genes passed all filters. **(B)**Workflow for deriving the gene indispensability index as a measure of the survival disadvantage a tumor cell incurs when it downregulates a specific gene.

From the 22 prioritized genes’ protein sequences, we proceeded to enumerate all MHC-I-restricted peptides whose sequence does not appear elsewhere in the human proteome. We then took advantage of published databases of MHC-peptide interactions and T-cell reactivity (MHCBN) to build machine learning-based predictors of MHC binding and immunogenicity (i.e., activity) derived from the peptides’ physicochemical properties. From prior knowledge, it is clear that the above naïve peptide enumeration generates a training set with a large majority of non-binders due to inclusion of ineligible peptides that would be discarded during the process leading up to MHC loading. The predictors’ performance was customized to handle this imbalance in dataset composition (see Methods). By maximizing positive predictive value and specificity **(Figure 3B),**we aimed to actively exclude false positives from the result list.

**Figure 3.**
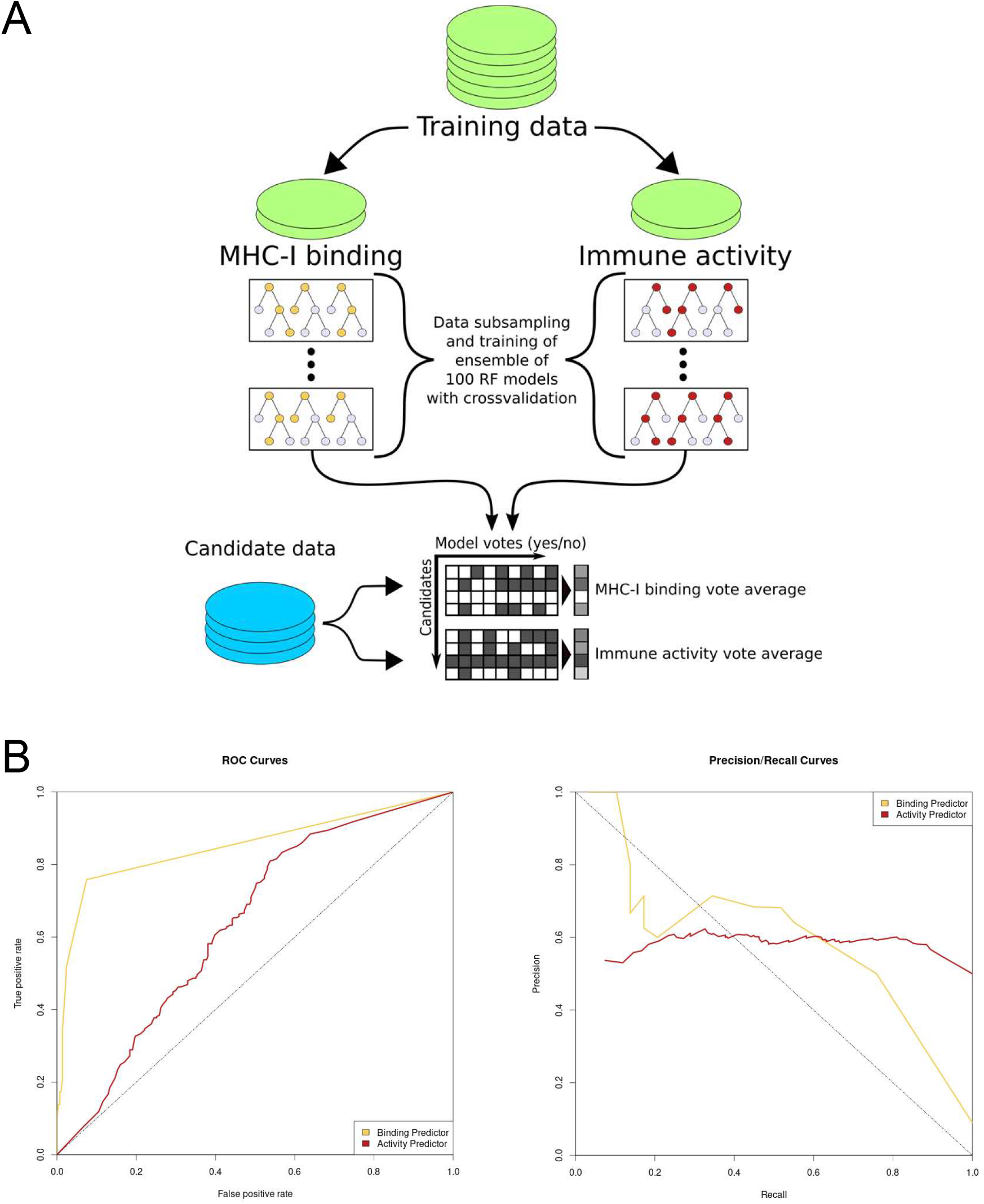
Training approach and performance of the dedicated machine learning predictors of MHC binding and T-cell receptor engagement (Immune activity). **(A)**For each of the two ensemble models, the training data was subsampled 100 times at a target-to-control sample ratio of 1:10 (binding) or 1:1 (activity) and used to construct 100 random forest (RF) models with 10,000 trees each. Weighted sampling was performed to reproduce the empirically observed surplus of non-binding peptides. **(B)**Receiver operating characteristics (ROC) and precisionrecall plots for the binding and the activity model. Area under the ROC curve (AUC) for the binding model was 0.86 while AUC for the activity model was 0.65.

**Figure 4.**
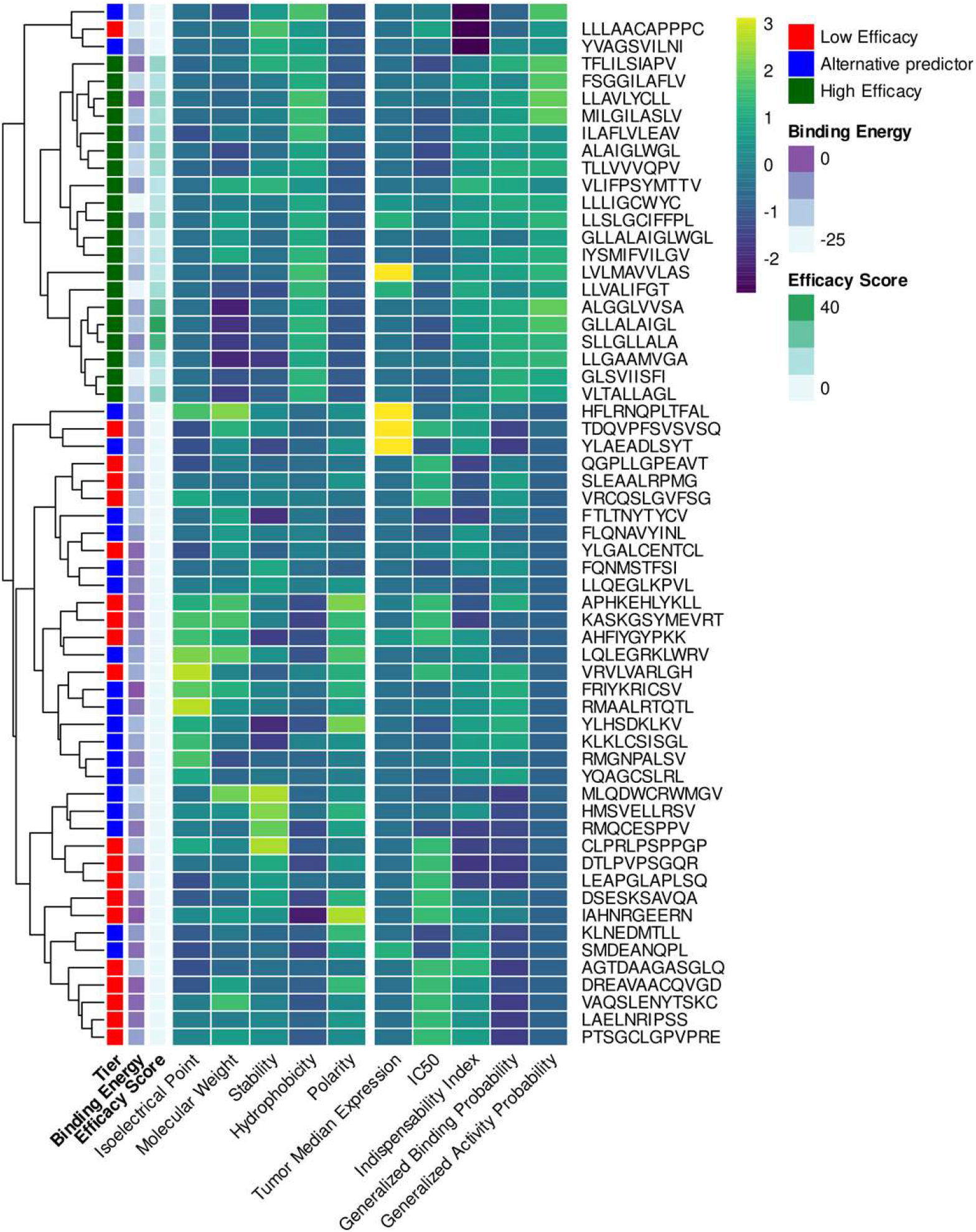
Heat map visualizing patterns in the factors contributing to the efficacy scores of the 60 selected peptide candidates. The columns show the physicochemical peptide features used to train the binding and activity predictors (left and the factors in the efficacy score (ES) equation (right) after z-score transformation. The IC50 column holds the peptides’ netMHCpan-predicted binding affinity to MHC for the HLA-A*02:01 allele. Rows are labeled with the peptide’s amino acid sequence and annotated with the ES, the computationally calculated binding energy to MHC (A*02:01), and the allocated ES tier. The high-efficacy group peptides are characterized by high hydrophobicity, an observation that is in line with established knowledge 1,2.

### Peptide ranking according to predicted efficacy

Antigenic peptide efficacy was calculated by plugging the obtained results into **equation III.**The product structure of the right-hand formula is modeled as a chained probability, with every factor representing a probability of success. Consequently, the efficacy scores fall within the interval between 0 and 1. Comparing the 60 efficacy score values against their constituent factors and the corresponding peptide properties, we observed no linear dependence, which rules out the possibility that the efficacy score is dominated by one or two of the constituents while the others contribute nothing.

The tumor expression consTME(gene(P)) and binding affinity consIC50(E) were integrated with the following rationale: With the first, we recover the median expression in the tumor as a measure of antigen availability for different genes. With the second, we complement our own binding predictor with the results of netMHCpan, in part to recover MHC allele specificity and also to enable the validation strategy described in the next section.

## Discussion

Metastatic uveal melanoma (UM) is a cancer with a bleak prognosis. Aiming to discover additional therapy options, we provide and validate an antigen selection algorithm for targeted therapies like autologous T-Cell transfer or therapeutic antitumor vaccination. Our approach for MHC-I-restricted antigens explicitly optimizes for self-tolerance and anti-tumor immunogenicity at the selection level. While there are many MHC-I binding prediction algorithms published that focus on the interaction of peptide and HLA-Allele, we show that a tumor entity-specific approach incorporating the relevant transcriptomic landscapes can yield promising results. In particular, our efficacy score and validation pipeline can be extended to other cancer entities for which transcriptomic information is available with little need for adaptation.

Compared to other antigen and peptide selection algorithms, ours starts from expression data without prior knowledge but integrates two mechanisms to avoid potentially life-threatening autoimmune reactions in non-tumor tissues: discarding genes with high expression in critical tissues and discarding peptides showing coincidental sequence identity with non-selected antigens.

The methodology presented in this study lends itself to the design of targeted anti-UM immunotherapies in both the non-personalized and the personalized setting. With contemporary turn-around times in RNA-sequencing and GMP-grade peptide synthesis, an individual patient’s tumor biopsy transcriptome can inform therapeutic decisions with only marginal delay. Beyond that, however, large cohort studies of the UM transcriptome will potentially allow the identification of antigens and peptides that are simultaneously efficacious and tolerable in a high percentage of UM patients. This, in turn, will make it possible to generate pre-manufactured libraries of peptides that are immediately available for use and bring the vision of off-the-shelf antigen therapies closer to reality.

## Funding statement

Manfred-Roth-Stiftung and the intramural Forschungsstiftung Medizin at the Universitätsklinkum Erlangen in a project on epitope prediction for uveal melanoma [to J.V.]. Hiege-Stiftung in a project on CAR T-cell development [to J.V.]. German Ministry of Education and Research (BMBF) in projects e:Med MelAutim [01ZX1905A to J.V.] and Kl-VesD [161L0244A to J.V.].

## Conflict of interest

The authors declare no conflict of interest related to the content of the paper.

## Notes

### Competing Interest Statement

The authors have declared no competing interest.

