## Supplementary Material for "An algorithm that combines machine learning ensemble modeling and network analysis to predict self-tolerant tumor-associated antigens for anti-cancer immunotherapy"

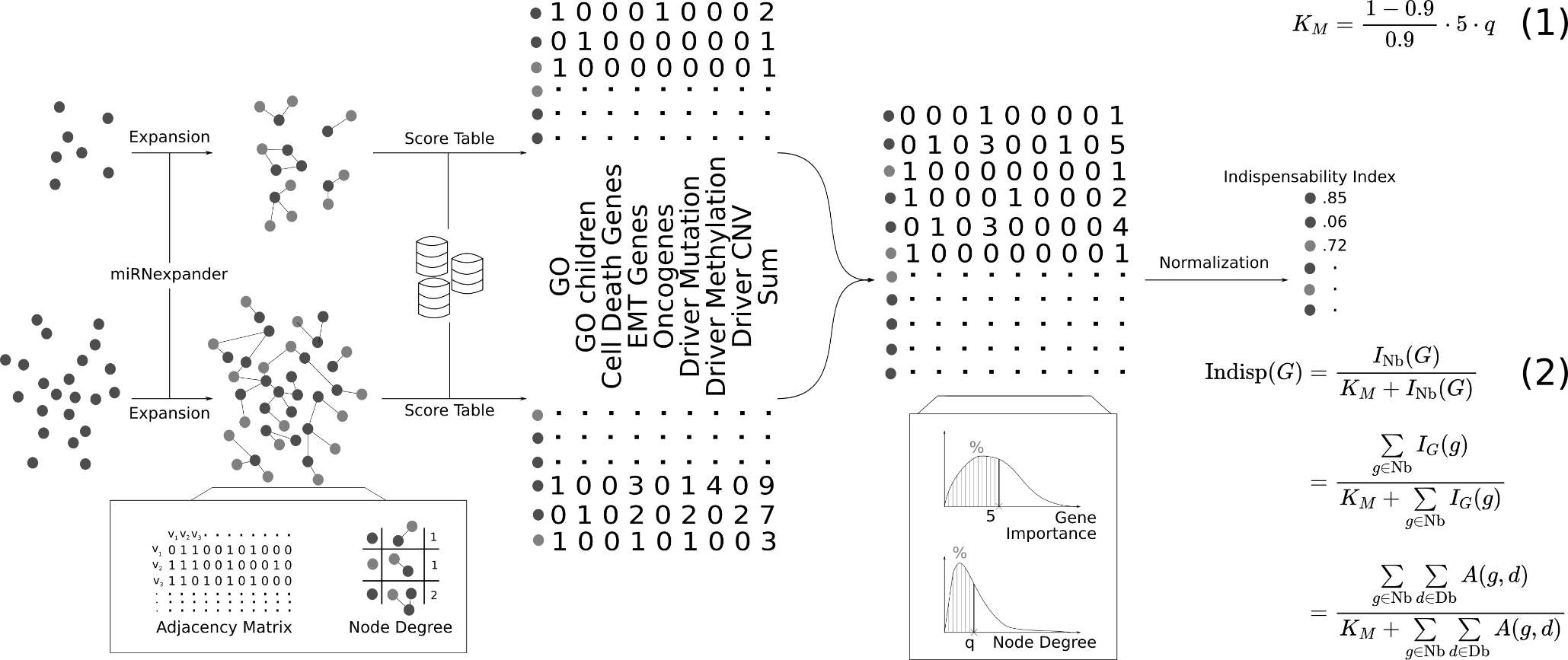

**Supplementary Figure S1. Further details on the derivation of the gene indispensability index.** This figure is a more technical representation of Figure 2C. The predicted genes and all the genes contained in DriverDBv3 are expanded into interaction networks via miRNexpander. For the expanded DriverDBv3 network, its adjacency matrix and the node degrees are calculated. For each network, node scores are tabulated by counting the occurrence of each node’s corresponding gene in several databases (see Methods). The sum of all occurrences becomes a gene’s importance. The two score tables are merged along the row dimension into one matrix. From the distributions of gene importance and node degree, normalization parameters for the next step are extracted. By multiplying the adjacency matrix and the sum column of the score matrix, each gene obtains a novel value termed neighborhood importance. The neighborhood importance is transformed into a variable with saturation characteristics via a Michaelis-Menten type equation (Eq. 2), with K_M_ calculated from the previously derived normalization parameters of gene importance and node degree (Eq. 1).

In **equation 1**, q and the number 5 denote the quantiles of gene importance and node degree, respectively, which were selected to estimate K_M_.

In **equation 2**, Indisp is the indispensability index function, I_Nb_ is the neighborhood importance function, and I_G_ denotes the gene importance function. G denotes a gene of interest, Nb the neighborhood around it, and g one gene belonging to G’s neighborhood, e.g. all of G’s immediate neighbors and G itself. Db denotes the set of databases (GO terms and cancer-related) on which the gene importance is based, d a member of Db, and A the function that returns the number of associations between database d and gene g.

**Supplementary Table S1: Manually curated cancer-associated GO Terms.** Genes associated with the terms in this list were scored higher during the indispensability index computation. Terms in bold font were selected by hand while the other terms were included due to being their descendants in the GO Term hierarchy.

| **GOID** | **Term** |
| --- | --- |
| **GO:0001525** | **angiogenesis** |
| GO:0001569 | branching involved in blood vessel morphogenesis |
| GO:0001833 | inner cell mass cell proliferation |
| GO:0001834 | trophectodermal cell proliferation |
| **GO:0001837** | **epithelial to mesenchymal transition** |
| GO:0002040 | sprouting angiogenesis |
| GO:0002041 | intussusceptive angiogenesis |
| GO:0002174 | mammary stem cell proliferation |
| GO:0002674 | negative regulation of acute inflammatory response |
| GO:0002677 | negative regulation of chronic inflammatory response |
| GO:0002862 | negative regulation of inflammatory response to antigenic stimulus |
| GO:0002941 | synoviocyte proliferation |
| GO:0003347 | epicardial cell to mesenchymal cell transition |
| GO:0003419 | growth plate cartilage chondrocyte proliferation |
| **GO:0006915** | **apoptotic process** |
| GO:0006925 | inflammatory cell apoptotic process |
| **GO:0008283** | **cell proliferation** |
| GO:0008284 | positive regulation of cell population proliferation |
| GO:0008285 | negative regulation of cell population proliferation |
| GO:0008637 | apoptotic mitochondrial changes |
| GO:0010463 | mesenchymal cell proliferation |
| GO:0010657 | muscle cell apoptotic process |
| GO:0010717 | regulation of epithelial to mesenchymal transition |
| GO:0010718 | positive regulation of epithelial to mesenchymal transition |
| GO:0010719 | negative regulation of epithelial to mesenchymal transition |
| GO:0014009 | glial cell proliferation |
| GO:0014029 | neural crest formation |
| GO:0016525 | negative regulation of angiogenesis |
| GO:0033002 | muscle cell proliferation |
| GO:0033028 | myeloid cell apoptotic process |
| GO:0033687 | osteoblast proliferation |
| GO:0034349 | glial cell apoptotic process |
| GO:0035172 | hemocyte proliferation |
| GO:0035492 | negative regulation of leukotriene production involved in inflammatory response |
| GO:0035726 | common myeloid progenitor cell proliferation |
| GO:0035736 | cell proliferation involved in compound eye morphogenesis |
| GO:0035988 | chondrocyte proliferation |
| GO:0036093 | germ cell proliferation |
| GO:0042127 | regulation of cell population proliferation |
| GO:0042981 | regulation of apoptotic process |
| GO:0043065 | positive regulation of apoptotic process |
| GO:0043066 | negative regulation of apoptotic process |
| GO:0043276 | anoikis |
| GO:0044340 | canonical Wnt signaling pathway involved in regulation of cell proliferation |
| GO:0044346 | fibroblast apoptotic process |
| GO:0045765 | regulation of angiogenesis |
| GO:0045766 | positive regulation of angiogenesis |
| GO:0048134 | germ-line cyst formation |
| GO:0048144 | fibroblast proliferation |
| GO:0050673 | epithelial cell proliferation |
| **GO:0050728** | **anti-inflammatory response** |
| GO:0051402 | neuron apoptotic process |
| GO:0051450 | myoblast proliferation |
| GO:0060055 | angiogenesis involved in wound healing |
| GO:0060266 | negative regulation of respiratory burst involved in inflammatory response |
| GO:0060317 | cardiac epithelial to mesenchymal transition |
| GO:0060722 | cell proliferation involved in embryonic placenta development |
| GO:0060809 | mesodermal to mesenchymal transition involved in gastrulation |
| GO:0060886 | clearance of cells from fusion plate by epithelial to mesenchymal transition |
| GO:0060978 | angiogenesis involved in coronary vascular morphogenesis |
| GO:0061323 | cell proliferation involved in heart morphogenesis |
| GO:0061351 | neural precursor cell proliferation |
| GO:0070341 | fat cell proliferation |
| GO:0070661 | leukocyte proliferation |
| GO:0071335 | hair follicle cell proliferation |
| GO:0071838 | cell proliferation in bone marrow |
| GO:0071839 | apoptotic process in bone marrow cell |
| GO:0071887 | leukocyte apoptotic process |
| GO:0072089 | stem cell proliferation |
| GO:0072104 | glomerular capillary formation |
| GO:0072111 | cell proliferation involved in kidney development |
| GO:0090255 | cell proliferation involved in imaginal disc-derived wing morphogenesis |
| GO:0097152 | mesenchymal cell apoptotic process |
| GO:0097190 | apoptotic signaling pathway |
| GO:0097194 | execution phase of apoptosis |
| GO:0097360 | chorionic trophoblast cell proliferation |
| GO:0106015 | negative regulation of inflammatory response to wounding |
| GO:0140208 | apoptotic process in response to mitochondrial fragmentation |
| GO:0150079 | negative regulation of neuroinflammatory response |
| GO:1900016 | negative regulation of cytokine production involved in inflammatory response |
| GO:1902362 | melanocyte apoptotic process |
| GO:1902489 | hepatoblast apoptotic process |
| GO:1902742 | apoptotic process involved in development |
| GO:1903594 | negative regulation of histamine secretion by mast cell |
| GO:1904019 | epithelial cell apoptotic process |
| GO:1904516 | myofibroblast cell apoptotic process |
| GO:1904606 | fat cell apoptotic process |
| GO:1990009 | retinal cell apoptotic process |
| GO:1990654 | sebum secreting cell proliferation |
| GO:2000793 | cell proliferation involved in heart valve development |

**Supplementary Table S2: HLA alleles taken into account when running the prediction pipeline.** These 36 alleles are the ones netMHCpan is pre-trained for. A*02:01 (in bold) is the allele for which peptides for validation experiments were selected. The other 35 alleles are in this context referred to as “bystander” alleles.

| **HLA Alleles** | | |
| --- | --- | --- |
| A*01:01 | B*07:02 | C*01:02 |
| **A*02:01** | B*08:01 | C*02:02 |
| A*03:01 | B*13:02 | C*04:01 |
| A*11:01 | B*15:01 | C*05:01 |
| A*24:02 | B*18:01 | C*06:02 |
| A*25:01 | B*27:02 | C*07:01 |
| A*26:01 | B*35:01 | C*08:02 |
| A*29:02 | B*41:01 | C*12:03 |
| A*31:01 | B*44:02 | C*15:02 |
| A*33:01 | B*44:03 | C*16:01 |
| A*68:01 | B*49:01 |  |
|  | B*50:01 |  |
|  | B*51:01 |  |
|  | B*52:01 |  |
|  | B*57:01 |  |
